# Respiration-driven brain network: neural underpinning of breathing correlates with resting-state fMRI signal

**DOI:** 10.1101/2022.07.11.499588

**Authors:** Wenyu Tu, Nanyin Zhang

## Abstract

Respiration can induce motion and CO_2_ fluctuation during resting-state fMRI (rsfMRI) scans, which will lead to non-neural artifacts in the rsfMRI signal. In the meantime, as a crucial physiologic process, respiration that can directly drive neural activity change in the brain, and may thereby modulate the rsfMRI signal. Nonetheless, this potential neural component in the respiration-fMRI relationship is largely unexplored. To elucidate this issue, here we simultaneously recorded the electrophysiology, rsfMRI and respiration signals in rats. Our data show that respiration is indeed associated with neural activity changes, evidenced by a phase-locking relationship between slow respiration variations and the gamma-band power of the electrophysiologic signal recorded in the anterior cingulate cortex. Intriguingly, slow respiration variations are also linked to a characteristic rsfMRI network, which is mediated by gamma-band neural activity. In addition, this respiration-related brain network disappears when brain-wide neural activity is silenced at an iso-electrical state, while the respiration is maintained, further confirming the necessary role of neural activity. Taken together, this study identifies a respiration-related brain network underpinned by neural activity, which represents a novel component in the respiration-rsfMRI relationship that is distinct from respiration-related rsfMRI artifacts. It opens a new avenue for investigating the interactions between respiration, neural activity and resting-state brain networks in both healthy and diseased conditions.

## Introduction

Resting state fMRI (rsfMRI), which measures spontaneous blood-oxygen-level dependent (BOLD) signal, is a powerful tool for non-invasively investigating brain-wide functional connectivity ^1–3^. Due to its hemodynamic nature, the rsfMRI signal is susceptible to systemic physiological changes such as respiration and cardiac pulsations ^4–7^, and these effects are usually treated as non-neuronal artifacts in rsfMRI data.

Respiration is a major physiological process that drives fluctuations in cerebral blood flow and oxygenation ^5^. Respiration can affect the BOLD signal ^8–10^ with two types of effects resulting from slow respiration variations and fast cyclic changes, respectively. Slowly varying changes (typically below 0.15 Hz) in breathing rate and depth, which can be quantified as respiration volume per time (RVT) ^10^, covary with arterial CO_2_ ^11–14^ and affect the BOLD signal via vasodilatory effects and/or autonomic influences of the vessel tone ^15,16^. On the other hand, faster respiratory cycles (i.e. periodic inspiration and expiration), accompanied by the corresponding chest and neck movement, can induce changes in the static magnetic field, which in turn lead to BOLD signal changes ^17–19^. Both effects can be effectively mitigated/removed by standard rsfMRI preprocessing methods ^10,17^.

In addition to the well-characterized respiration-related non-neural artifacts, there is evidence hinting a possible neural component in the respiration-rsfMRI interaction ^20,21^. First, respiration can directly drive brain-wide neuronal oscillations ^22,23^ not only in the olfactory bulb ^24^ and piriform cortex ^25^—brain regions directly related to breathing— but also in the medial prefrontal cortex (mPFC), somatosensory cortex and hippocampus, and this effect has been consistently found across species ^26–29^. In addition, respiration changes can be associated with arousal and/or emotion-related brain state changes, which covary with cortical activity ^30–33^. Therefore, in addition to the artifactual effects aforementioned, respiration may affect the rsfMRI signal by directly modulating the neural activity. However, this potential neural component in the respiration-fMRI relationship is largely unexplored.

To gain a comprehensive understanding of the relationships between respiration, neuronal activity and rsfMRI signal, here we simultaneously acquired rsfMRI, electrophysiology and respiration data in anesthetized rats. Anesthesia was used to ensure our results are not confounded by the animal’s motion, which affects all three signals. Based on these measures, an RVT-correlated rsfMRI network was identified. Importantly, regressing out gamma activity or silencing neural activity across the brain disrupted this respiration-related network, suggesting that this respiration-rsfMRI relationship is mediated by neural activity.

## Results

To determine the potential role of neural activity in the respiration-rsfMRI relationship, we simultaneously recorded the electrophysiology and respiration signals along with rsfMRI data in rats (Fig. 1A). The respiration signal was recorded by a respiration sensor placed under the animal’s chest (Fig. 1A). Representative raw respiration signal, the respiration rate distribution across all scans, and the averaged respiration power are shown in Fig. 1B&C. Slow respiration variations are quantified by RVT, calculated as the difference of consecutive peaks of inspiration and expiration divided by the time interval between the two adjacent signal maxima (or minima) (Fig. 1D) ^10^. The electrophysiology signal was recorded using an MR-compatible electrode implanted in the right side of the ACC. The ACC was selected given its critical role in respiratory control ^34^. In addition, the ACC is a key region in the rodent default-mode network (DMN) ^35,36^, and the DMN has been linked to respiration-related fMRI signal changes ^10^. The location of the electrode was confirmed by T2-weighted structural images (Figure 1 – Figure Supplement 1). MRI artifacts in the electrophysiology signal was removed using a template regression method ^37,38^ (see Methods and Materials and Fig.S6), and local field potential (LFP) and spectrogram were obtained using denoised electrophysiologic data (Fig. 1E).

**Figure 1.**
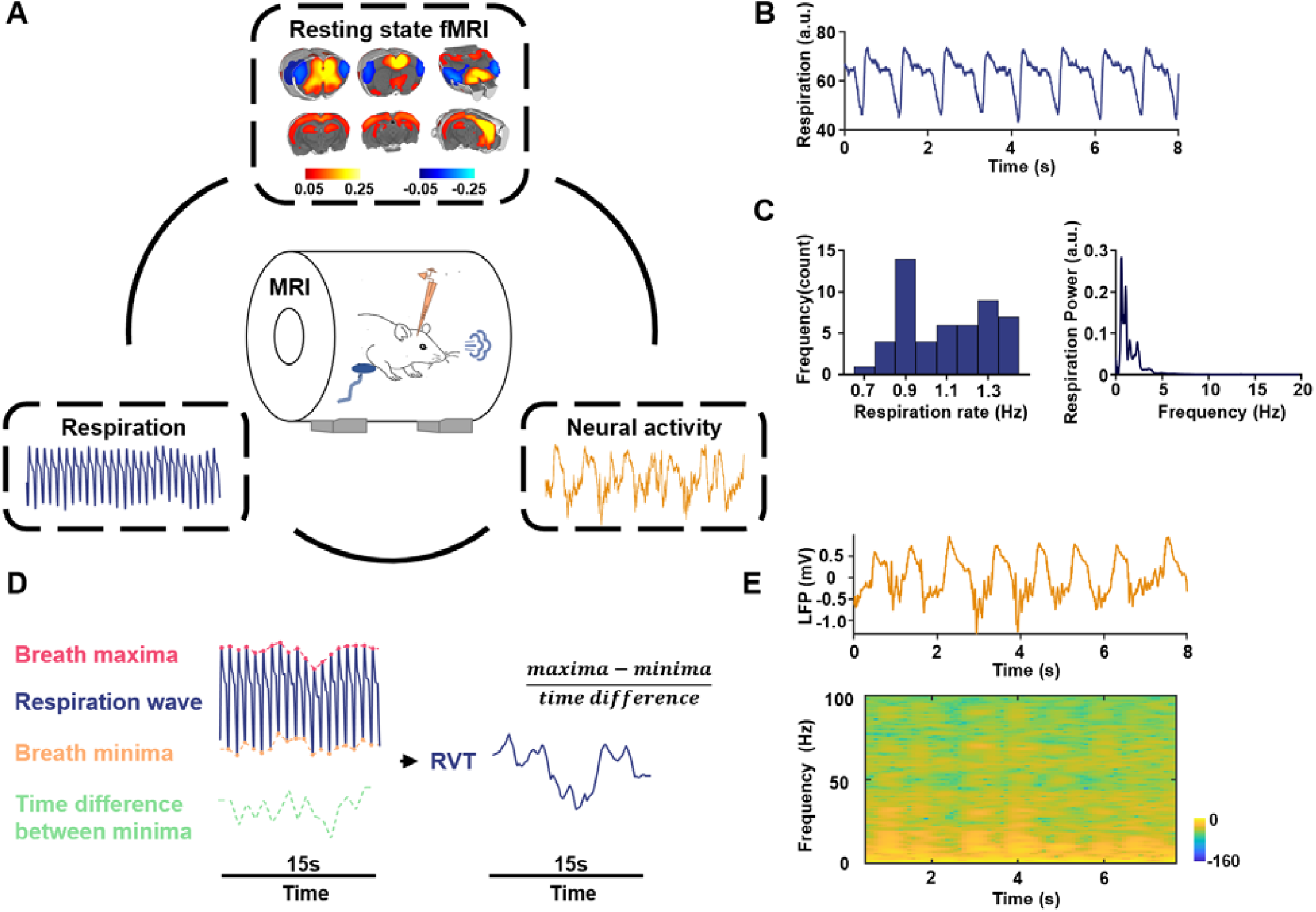
A. Experimental Design – simultaneous measurement of the rsfMRI, electrophysiology and respirational signals. Top: ACC seedmap; bottom left: respiration signal; bottom right: local field potential. **B**. Exemplar respiration signal waveform. **C**. Left: distribution of the respiration rate across all scans; right: power of the respiration signal averaged across all scans. **D**. Computing RVT from the respiration waveform. **E**. Exemplar denoised LFP signal. Top: LFP time series; bottom: LFP power spectrogram.

### Gamma-band neural oscillations are respectively associated with respiration and rsfMRI signals

We first asked whether the respiration is linked to the neural activity changes (Fig. 2A) in lightly sedated animals (combined low-dose dexmedetomidine and isoflurane, see Methods and Materials). Prominent coherence between the respiration signal and LFP is observed with the dominant peak at ~1.1 Hz (Fig. 2B&C), in line with the respiration frequency in animals (Fig. 1C). This result is consistent with the finding of respiration entrained LFP oscillations previously reported ^22,23^.

**Figure 2.**
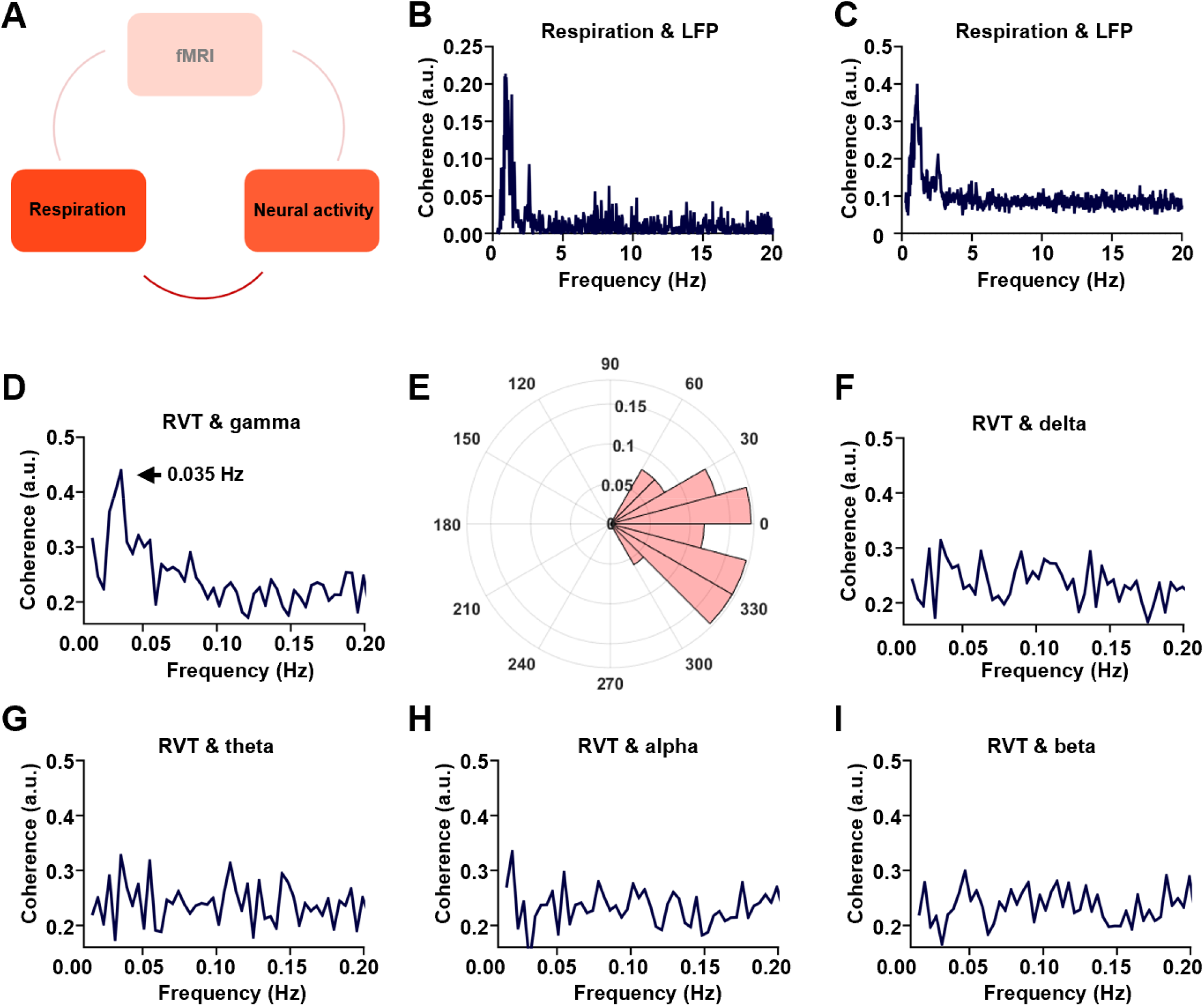
Phase locking relationship between slow respiration variations and neural activity. **A**. The relationship between respiration and neural activity. **B**. Respiration-LFP coherence from one sample scan. **C**. Respiration-LPF coherence averaged across all scans. **D**. Coherence between RVT and gamma band power (40-100 Hz), with the peak at 0.035 Hz. **E**. Phase lag between RVT and gamma band power. In contrast, no obvious coherence is observed between RVT and delta band power (1-4 Hz, **F**), theta band power (4-7 Hz, **G**), alpha band power (7-13 Hz, **H**), or beta band power (13-30 Hz, **I**).

To further dissect the respiration-LFP relationship, the LFP was separated into five conventionally defined frequency bands including gamma (40-100 Hz), delta (1-4 Hz), theta (4-7 Hz), alpha (7-13 Hz) and beta band (13-30 Hz) ^39,40^. Gamma band power displays significant coherence with the RVT at ~0.035 Hz with a zero phase lag (Fig. 2D&E, phase (mean ± ste) = −0.08±0.5 pi, with the range of [-pi, pi]), suggesting a phase locking relationship between the two measures. In contrast, this coherence is not observed in any other frequency bands (Fig. 2 F-I). These results remain the same when the LFP data were analyzed using the local subtraction method (Figure 2 – Figure Supplement 1A-C), ruling out the potential artifact resulting from the volume conduction of signals between the cortex and olfactory bulb ^41^. Taken together, our data demonstrate that respiration is associated with gamma-band neural oscillations in the ACC.

We next examined the relationship between the gamma-band LFP and rsfMRI signals (Fig. 3A) ^42^. The ACC gamma power was first convolved with a hemodynamic response function (HRF, defined by a single gamma probability distribution function (a = 3, b = 0.8) ^43^, Fig. 3B), which was then voxel-wise correlated to brain-wide rsfMRI signals. Our data show that the gamma power-derived correlation map (Fig. 3C) is highly consistent with the ACC RSFC seedmap with the right ACC (i.e. the electrode implanted side) as the seed (Fig. 3D), evidenced by a strong voxel-to-voxel spatial correlation between these two maps (Fig. 3E, R = 0.775, p < 10^−15^). Again, the same gamma power-derived correlation pattern is observed when the LFP data were analyzed using the local subtraction method (Figure 2 – Figure Supplement 1D). These data demonstrate that the gamma-band LFP is tightly linked to the rsfMRI signal.

**Figure 3.**
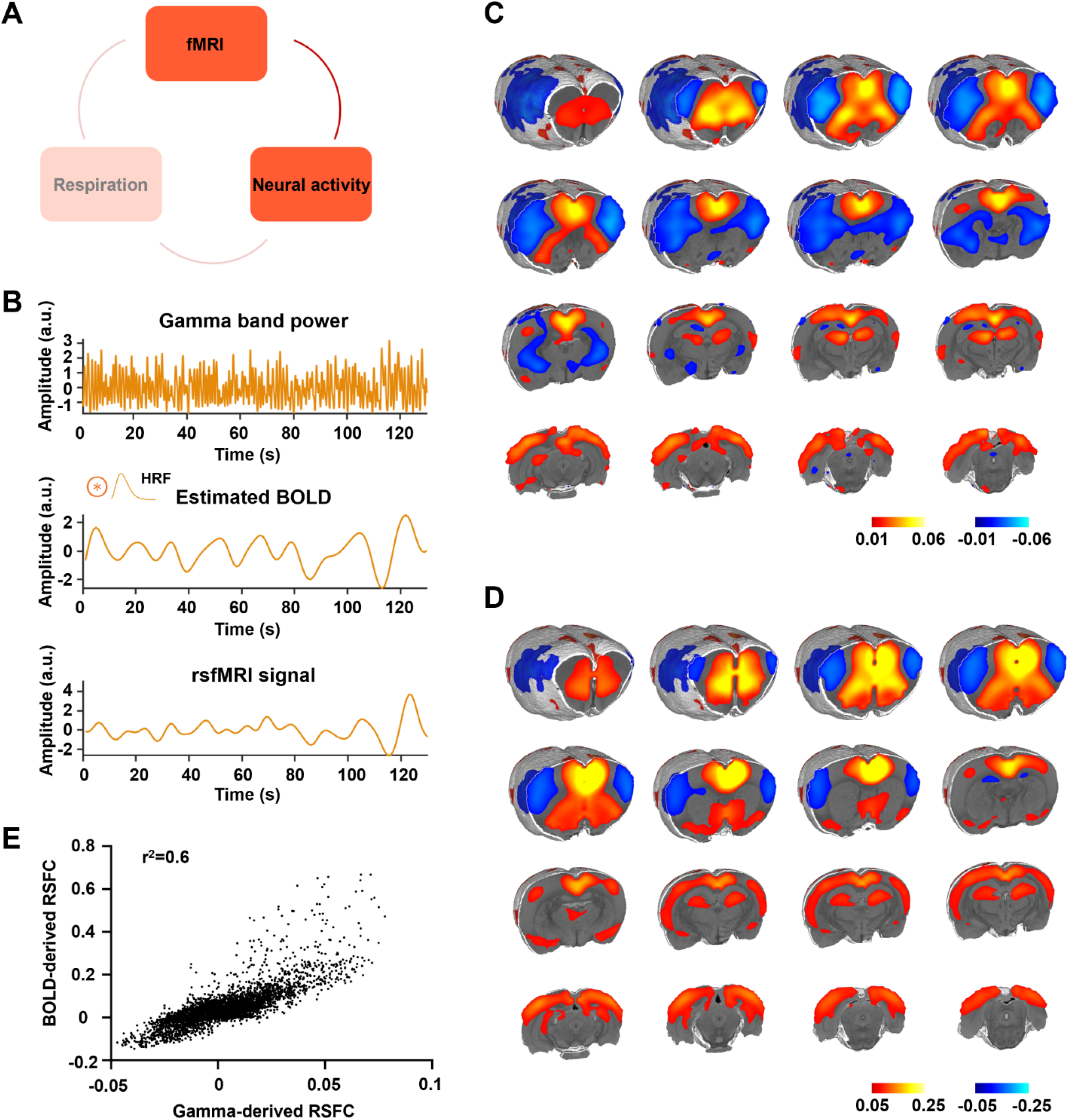
Gamma power is associated with the rsfMRI signal. **A**. The relationship between neural activity and rsfMRI signal. **B**. Top: exemplar gamma band power in the ACC; middle: estimated BOLD signal by convolving the gamma band power with the hemodynamic response function (HRF); bottom: measured BOLD signal from the same brain region. **C**. Gamma power-derived correlation map. **D**. Seedmap of the right ACC. **E**. Voxel-to-voxel spatial correlation between (C) and (D).

### Respiration drives a characteristic rsfMRI network, mediated by gamma-band neural activity

Given that the gamma band power is respectively associated with the RVT and rsfMRI signals, we specifically asked how RVT is related to the rsfMRI signal in lightly sedated animals by calculating voxel-wise correlations between the RVT and rsfMRI signals (Fig. 4A). This analysis generates a respiration-related rsfMRI network (Fig. 4B), involving key brain regions controlling respiration such as the piriform cortex. It is known that the piriform cortex receives inputs from the olfactory bulb, and can be directly activated when the animal breathes. In addition, this network includes regions involved in the rodent DMN such as the ACC, mPFC, orbital, retrosplenial and primary somatosensory cortices, as well as the hippocampus ^35^ (Fig. 4C, one-sample t-test, p < 0.05, FDR corrected). The resemblance of this respiration-related network and DMN well agrees with the human literature that the physiological effects on rsfMRI data are colocalized with the DMN ^10^. Taken together, slow respiration variations exhibit a characteristic correlation pattern with brain-wide rsfMRI signals, representing a respiration-related rsfMRI network.

**Figure 4.**
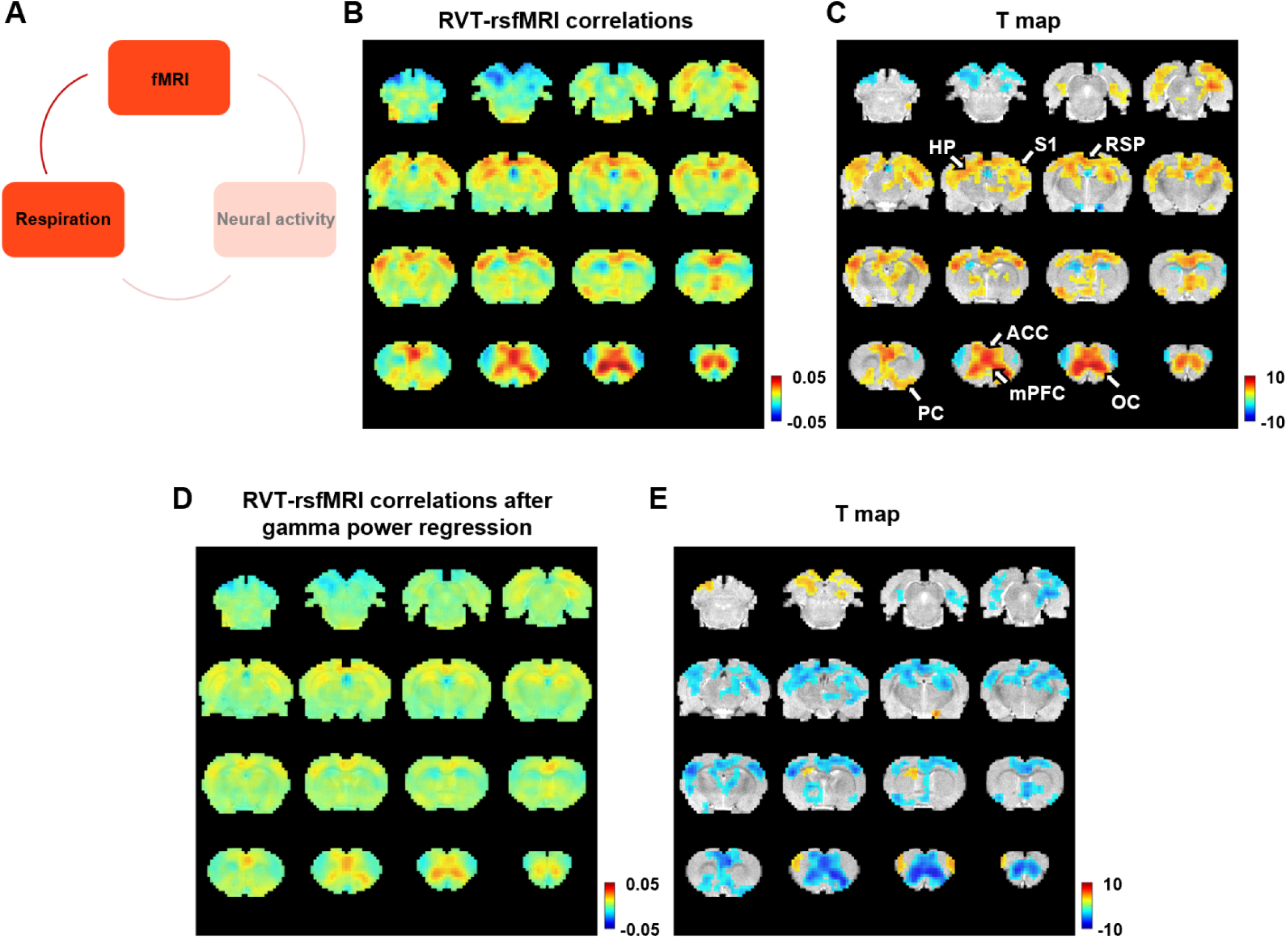
Correlation between slow variations of respiration and rsfMRI signal. **A**. The relationship between respiration and rsfMRI signals. **B-C**. Voxelwise correlations between the RVT and rsfMRI signals. **B**. Unthresholded correlation map averaged across scans. **C**. Thresholded T-value map (one-sample t test, p < 0.05, FDR corrected). Brain regions displaying significant RVT-rsfMRI correlations include the anterior cingulate cortex (ACC), orbital cortex (OC), medial prefrontal cortex (mPFC), piriform cortex (PC), hippocampus (HP), retrosplenial cortex (RSP) and primary somatosensory cortex (S1). **D**. Voxelwise correlations between the RVT and rsfMRI signals after the gamma-band power is regressed out from both signals. **E**. Difference of correlation maps before and after gamma power regression (paired t-test, p < 0.05, FDR corrected).

To test whether this respiration-related rsfMRI network is mediated by gamma-band activity, the gamma power was respectively regressed out from the RVT and voxelwise rsfMRI signals, and the same correlational analysis was repeated on the residual signals. In this case, the respiration-related network is diminished (Fig. 4D). This result is confirmed by the contrast of the correlation maps before (Fig. 4B) and after (Fig. 4D) gamma power regression, with essentially all brain regions involved in the respiration-related network exhibiting reduced RVT-rsfMRI correlation after the gamma power is regressed out (Fig. 4E, paired t-test, p < 0.05, FDR corrected). These results demonstrate that the respiration-related rsfMRI network is mediated by gamma-band neural activity.

### The respiration-related rsfMRI network is absent at the isoelectric state

To further confirm the necessary role of neural activity in the respiration-related rsfMRI network, we experimentally silenced the neural activity in the whole brain by inducing an isoelectric brain state using high-dose sodium pentobarbital, while maintained the respiration in the rat (Fig. 5A) ^44^. At the isoelectric state, the LFP amplitude and power are close to zero, in remarkable contrast to the electrophysiology data recorded before the drug infusion (Fig. 5B). In addition, rsfMRI data recorded at the isoelectric state exhibit a flat, noise-like power distribution, unlike the characteristic 1/f pattern during light sedation (Fig. 5D) ^45^. Furthermore, there is no meaningful ACC RSFC in the ACC seedmap at the isoelectric state (Fig. 5E), which is distinct from the ACC seedmap during light sedation (Fig. 3D). The brain-wide ROI-based RSFC matrix also reveals a global suppression of RSFC at the isoelectric state (Fig. 5G), compared to the RSFC matrix observed at the light sedation condition.

**Figure 5.**
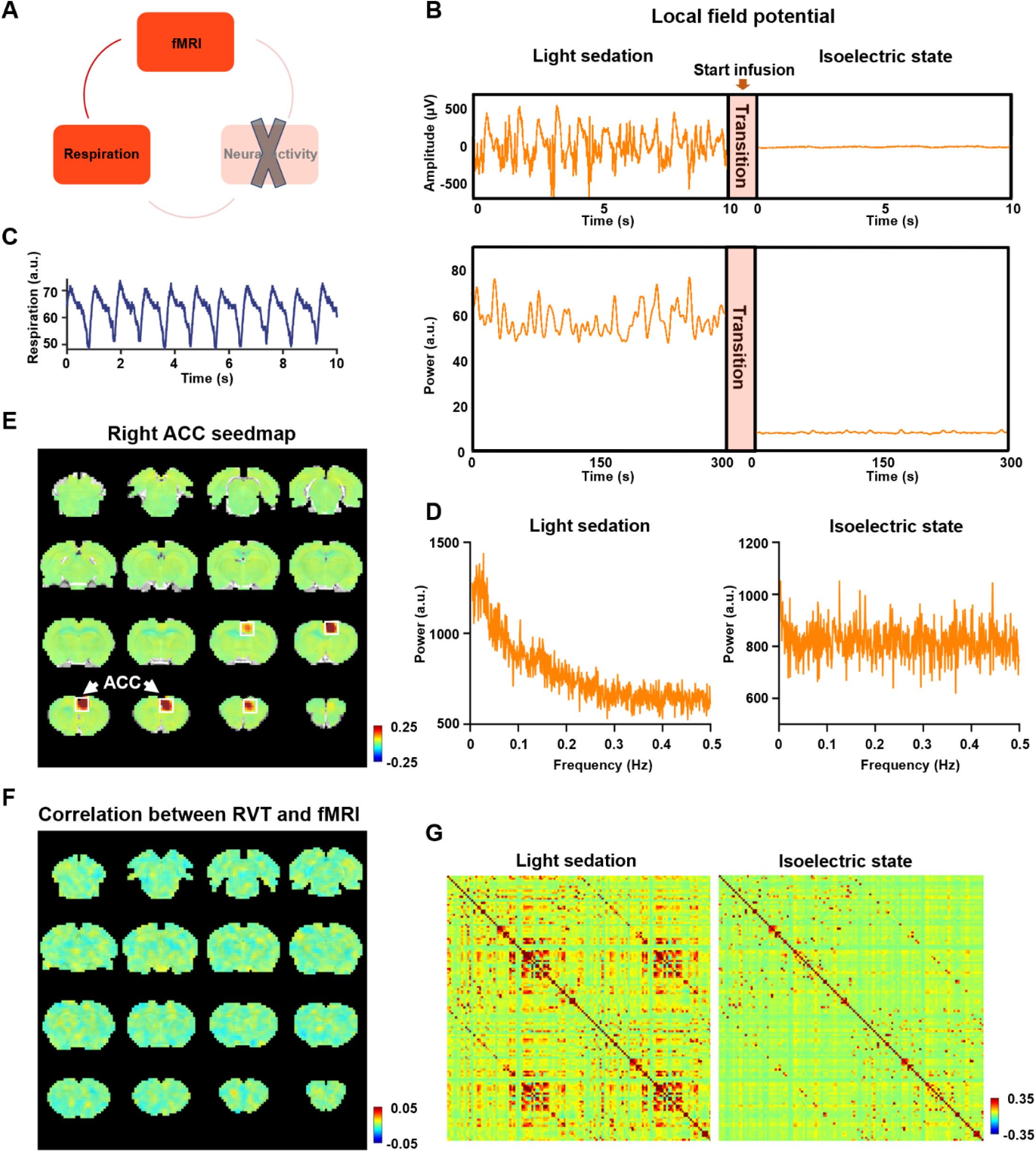
Respiration-rsfMRI relationship at the isoelectric state. **A**. Determining the relationship between slow respiration variations and the rsfMRI signal after silencing the brain-wide neural activity. **B**. Silencing neural activity at the isoelectric state induced by sodium pentobarbital. Top: LPF amplitude; bottom: LPF power; arrow: infusion of sodium pentobarbital. **C**. Respiration signal at the isoelectric state. **D**. Power spectra of the BOLD signal at (left) light sedation and (right) the isoelectric state. **E**. Seedmap of the right ACC at the isoelectric state. **F**. Voxelwise correlations between RVT and the rsfMRI signal at the isoelectric state. **G**. Brainwide ROI-based RSFC matrices at (left) light sedation and (right) the isoelectric state (132 ROIs in total).

After confirming the global neural silencing effect, we calculated voxel-wise correlations between the RVT and rsfMRI signals at the isoelectric state. The respiration-related network observed during light sedation is absent when brain-wide neural activity is silenced (Fig. 5F), despite similar breathing patterns (Fig. 5C) at both conditions. These data demonstrate that the neural activity plays a necessary role in the respiration-related rsfMRI network, corroborating the notion that the respiration-related rsfMR network we observed is mediated by neural activity.

### Respiration-related brain network is distinct from respiration-related rsfMRI artifacts

To confirm the respiration-driven neural network we observed is different from respiration-related rsfMRI artifacts previous reported, we separately obtained the artifactual patterns of fast (i.e. cyclic inspiration and expiration) and slow respiration signals in the rsfMRI data.

The fast variations of respiration can cause aliased respiratory artifacts in the fMRI signal by inducing B0 offsets ^46,47^. To determine this effect, we calculated voxel-wise correlations between the rsfMRI signal and the respiration regressor estimated by RETROICOR (See Methods for details) ^17^. Unlike the respiration-induced neural network described above, we did not observe any appreciable correlations between RETROICOR and the rsfMRI signals across the brain at either the light sedation or isoelectric state (Figure 4 – Figure Supplement 1). This result remains the same with and without regressing out the signals of the white matter and ventricles in rsfMRI preprocessing, likely due to minimal motion of the animal at both anesthetized states.

To assess the artifactual effects resulting from slow respiration signals on rsfMRI data, we adopted the standard analysis method by convolving the RVT with a respiration response function (RRF) determined by the difference of two gamma variate functions (See Methods and Materials, Figure 4 – Figure Supplement 2A) ^48^, and then voxel-wise correlated it to the rsfMRI signal (Figure 4 – Figure Supplement 2). Note that this calculation is different from the method we used to identify respiration-induced neural network, which directly correlated the RVT with the rsfMRI signal. Our data show that the slow respiration-related rsfMRI artifact is distinct from the respiration-induced neural network. First, the slow respiration-related rsfMRI artifact is dominantly located around the ventricles and large veins (Figure 4 – Figure Supplement 2B), whereas the respiration-induced neural network is mostly located at the gray matter with a distinct spatial pattern (Fig. 4B&C). In addition, the slow respiration-related rsfMRI artifact can be reduced by regressing out the signals in the white matter and ventricles (Figure 4 – Figure Supplement 2C), but the respiration-induced neural network persists after the regression. Taken together, these data indicate that the respiration-induced neural network we observed is not a non-neural physiological artifact but originates from neural activity.

## Discussion

In this study, we discovered a slow respiration variation-induced functional network in rodents. Importantly, we identified the neural underpinning of this network, indicated by the phase-locking relationship between the RVT and gamma band power in the ACC—a node in this network. Regressing out the gamma-band power disrupted the respiration network, suggesting a mediating role of neural activity in this network. To further validate this finding, we experimentally silenced the neural activity across the brain but maintained the respiration in the animal. In this condition, we again failed to observe the respiration-related brain network, confirming the neural underpinning of the network. Lastly, we showed that the respiration-driven neural network is distinct from respiration-related rsfMRI artifacts. Overall, our study maps brain-wide neural responses related to respiration. These data provide new insight into understanding the neural activity-mediated respiratory effects on resting state functional networks. Importantly, this breathing-related brain network might be altered in brain disorders, and thus our finding might potentially provide important clinical value.

### The respiration network is very likely linked to respiration-entrained brain-wide neural oscillations

Respiration-induced neuronal oscillations have been consistently observed across multiple species including rats, mice and humans ^27,29,49^. This phenomenon also seems to be robust in various physiological and behavioral states including freely moving awake and immobile awake conditions, as well as anesthetized states ^25,26,50^. Intriguingly, previous studies found that respiration specifically modulated the amplitude of gamma activity in the mPFC ^50–52^, consistent with our finding of band-specific coherence between the gamma power in the ACC and RVT. Notably, gamma power is generally believed to relate to the BOLD signal ^37,42^, which is also demonstrated in our study (Fig. 3). These results suggest that respiration can drive gamma oscillations, which further lead to BOLD signal changes in the respiration network.

In addition to the mPFC, previous work has shown that respiration-generated neuronal oscillations are phase locked to the breath rhythm across distributed brain regions including the olfactory bulb, primary piriform cortex, hippocampus and somatosensory barrel cortex ^25–28,53^. Those regions are highly consistent with brain regions in the respiration network we observed. Furthermore, the piriform cortex is anatomically connected to the mPFC, ACC and orbital cortex, all of which are also parts of the respiration network in our study ^54–57^. Taken together, the spatial pattern of the respiration network reported here well agrees with brain regions exhibiting neural oscillation changes driven by respiration.

Previous work also demonstrates that respiration-induced neuronal oscillations do not result from the movement of the muscle or electrode. For instance, the oscillations are different in the laminar amplitude along the hippocampus, with the maximal amplitude in the dentate gyrus (DG) ^27^. This finding well agrees with our data, displaying more prominent involvement of the DG than other parts of the hippocampus in the respiration network. In addition, our data show that only the gamma-band power, not other LFP bands, is coherent with slow variations of respiration, whereas the movement of electrode or muscle would lead to increased coherence across multiple frequency bands. Taken together, the respiration-related functional network we observed is very likely linked to the respiration-entrained gamma band oscillations in regions involved in the network.

Notably, regressing out the respirational signal does not abolish the RSFC measured by rsfMRI data, suggesting that the respiration does not account for all the effects of the RSFC measured. Figure 3 – Figure Supplement 1 shows the differences of ACC seedmaps and ACC gamma activity-derived correlation maps before and after regressing out the RVT. Despite that RVT regression reduced ACC RSFC particularly in brain regions involved in the respiration network as expected, the major RSFC pattern remains consistent, indicating that a dominant component of RSFC is not ascribed by the respiration effect.

### Respiration and arousal changes

It is possible that the neural contribution to the respiration-driven brain network is mediated by arousal changes, which is associated with both neural excitability and respiration ^30,31,58^. The cerebral blood flow can be regulated by the innervation from the basal forebrain and locus coeruleus, and both regions are directly related to arousal changes ^30,59,60^. Previous studies in humans demonstrated correlations among the fMRI signal, low-frequency fluctuations of respiration, and EEG alpha power ^20^. The fMRI-respiration correlate was stronger at the eye-closing resting state with a greater arousal-level change than the eye-open state. In addition, strong association was identified between alpha wave and alertness level. These data together suggest that the correlations between low-frequency fluctuations of respiration, alpha EEG power and the fMRI signal might be related to the fluctuation of wakefulness ^20^. However, we do not believe the arousal level plays a dominant role in our results, given that our study was performed in anesthetized animals. Accordingly, we did not observe any coherence between alpha band power and the slow variations of respiration, suggesting the respiration-driven fMRI network we observed must be attributed to a different mechanism than arousal.

## Methods and Materials

### Animals

The present study was approved by the Pennsylvania State University Institutional Animal Care and Use Committee (IACUC). Seven adult male Long-Evans rats (300-500g) were used. Rats were housed in Plexiglas cages with food and water provided ad libitum. The ambient temperature was controlled at 22-24°C under a 12h light :12h dark schedule.

### Surgery

Stereotaxic surgeries were performed to implant electrodes in animals for the electrophysiology recording. The rat was anesthetized with an injection of ketamine (40mg/kg) and xylazine (12mg/kg), and remained anesthetized throughout the surgery by 0.75% isoflurane delivered through an endotracheal catheter intubated (PhysioSuite, Kent Scientific Corporaition). Antibiotics baytril (2.5mg/kg) and long-acting analgesic drug buprenorphine were intramuscularly administered. During surgery, the temperature was monitored and maintained by a warming pad (PhysioSuite, Kent Scientific Corporaition). Heart rate and SpO_2_ were monitored with a pulse oximetry (MouseSTAT® Jr, Kent Scientific Corporation). An MR-compatible electrode (MRCM16LP, NeuroNexus Inc) was unilaterally implanted into the anterior cingulate cortex (ACC, coordinates: anterior/posterior +1.5, medial/lateral +0.5, dorsal/ventral - 2.8). The reference wire and grounding wire from the electrode were both connected to a silver wire placed in the cerebellum. After surgery, the animal was returned to the home cage and allowed to recover for at least one week.

### Simultaneous rsfMRI, respiration and electrophysiology recordings

rsfMRI experiments were performed on a 7T Bruker 70/30 BioSpec running ParaVision 6.0.1(Bruker, Billerica, MA) with a homemade single loop surface coil at the high field MRI facility at the Pennsylvania State University. T2*-weighted gradient-echo rsfMRI images were acquired using an echo planar imaging sequence with following parameters: repetition time (TR) = 1 s; echo time (TE) = 15 ms; matrix size = 64 × 64; field of view = 3.2 × 3.2 cm^2^; slice number = 20; slice thickness = 1mm; volume number = 1200. T2-weighted structural images were also obtained using a rapid acquisition with relaxation enhancement (RARE) sequence with the following parameters: TR = 3000 ms; TE = 40 ms; matrix size = 256 × 256; field of view = 3.2 × 3.2 cm^2^; slice number = 20; slice thickness = 1 mm; repetition number = 6.

During imaging, the animal was maintained at one of two anesthetized states: light sedation and iso-electric state. For imaging sessions under light sedation, animals were anesthetized with the combination of dexmedetomidine (initial bolus of 0.05 mg/kg followed by constant infusion at 0.1 mg·kg^−1^·h^−1^) and isoflurane (0.3%) ^61^, and spontaneous respiration was maintained in animals. For imaging sessions under the isoelectric state, sodium pentobarbital was administered with a 30 mg/kg bolus followed by continuous infusion (70 mg·kg^−1^·h^−1^) ^44^. Before imaging, the rat was intubated via tracheal, and the respiration was controlled by a ventilator (PhysioSuite, Kent Scientific Corporaition) throughout the entire imaging session. Three rats were in the light sedation group with a total of 51 scans and four rats were in the isoelectric state group with a total of 48 scans. During all imaging sessions, the temperature was measured by a rectal thermometer and maintained at 37°C with warm air. Eyes of the animal were protected from dryness using artificial tear.

During rsfMRI acquisition, the respiration signal was simultaneously recorded at the sampling rate of 225 Hz by a respiration sensor placed under the animal’s chest. Electrophysiology recording started 10 min before the beginning of rsfMRI acquisition and continued throughout the whole imaging session with a TDT recording system including an MR-compatible LP16CH headstage, PZ5 neurodigitizer amplifier, RZ2 BioAmp Processor and WS8 workstation (Tucker Davis Technologies Inc, Alachua, FL). The electrophysiology signal was sampled at 24414 Hz and the unfiltered raw signal was used for further data processing.

### Data preprocessing

rsfMRI data were preprocessed using a pipeline described in our previous publications ^43,62^. Briefly, rsfMRI images were first motion scrubbed based on relative framewise displacement (FD). Volumes with FD > 0.25mm and their adjacent preceding and following volumes were removed. Subsequently, data were preprocessed by performing motion correction (SPM12), co-registration to a defined atlas, spatial smoothing, as well as voxelwise nuisance regression of motion parameters and the signals of the white matter and ventricles.

Electrophysiology data were preprocessed to remove MR artifacts using a template regression method ^37,38^. Specifically, raw electrophysiology signal was first temporally aligned with the corresponding rsfMRI scan. The potential phase differences across 16 electrophysiology channels were corrected by calculating cross correlations of electrophysiology time series between channels, and the corrected signals from all 16 channels were summed. This summed signal was then segmented for each individual rsfMRI slice acquisition. Subsequently, an MRI interference template for each fMRI slice was estimated by averaging raw electrophysiology data across all segments corresponding the same slice acquisition from all rsfMRI volumes. The template was then aligned to each slice acquisition using cross correlation. The final templates of all slices were linearly regressed out from the raw electrophysiology data to remove MR-induced artefacts, followed by a series of notch filters for harmonics of the power supply (60Hz and multiples of 60Hz) and slice acquisition (20Hz and multiples of 20Hz). Lastly, the continuous LFP was bandpass filtered (0.1 - 300 Hz). Figure 1 – Figure Supplement 2 shows an example of raw electrophysiology signals before denoising of MRI artifacts (Figure 1 – Figure Supplement 2A), an example of the MRI artifact template (Figure 1 – Figure Supplement 2B), as well as the LFP signal after MRI artifact denoising (Figure 1 – Figure Supplement 2C).

The LFP spectrogram was computed using the MATLAB function *spectrogram* with window size = 1 s, step size = 0.1 s, as shown in fig. 1E.

### Data analysis

To determine the relationship between slow variations of respiration and the rsfMRI signal, Pearson correlation was voxel-wise calculated between the rsfMRI signal and RVT, which was calculated by the difference between consecutive peaks of inspiration and expiration divided by the time interval between the two peaks ^10^. One sample t test was performed to determine the statistical significance of correlations, thresholded at p < 0.05 after false discovery rate (FDR) correction of multiple comparisons.

To determine the artefactual impact of slow respiration variations on the rsfMRI signal, we convolved the RVT with the respiration response function (RRF), and then voxel-wise correlated it to the rsfMRI signal. The RRF is defined below:

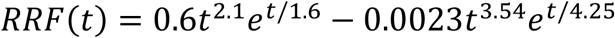

The fast cyclic respiratory effects on the rsfMRI signal were quantified using the method RETROICORR ^17^. Sinusoids were constructed based on TR relative to the phase of the respiratory cycles. For each brain voxel, the predicted fast respiration regressor was estimated by the linear combination of sinusoids that best fits to the voxel timeseries.

The phase locking relationship between the respiratory rhythm and the LFP oscillations was determined by calculating the magnitude squared coherence between the two signals. We also assessed the relationship between the frequency band-specific power of the LFP signal and slow variations of respiration. LFP band power was obtained using the MATLAB function *spectrogram* with window size = 1 s, step size = 0.1 s based on the conventional LFP band definition from previous studies (delta: 1-4 Hz, theta: 4-7 Hz, alpha: 7-13 Hz, beta: 13-30 Hz, gamma: 40-100 Hz) ^39,63^. The magnitude squared coherence and the phase relationship between the power of each LFP band and RVT was calculated using the MATLAB function *mscohere*.

To determine the relationship between the gamma band power and rsfMRI signal, the time course of gamma power was first convolved with the hemodynamic response function (HRF, a single gamma probability distribution function (a = 3, b = 0.8) ^43^), and voxel-wise Pearson correlations between the convolved gamma power and rsfMRI signals in the brain were computed.

The ACC seedmap was obtained by voxel-wise calculating the Pearson correlations between the regionally averaged rsfMRI time course of the unilateral right-side ACC with the rsfMRI signals across the whole brain. To perform the region of interest (ROI)-based analysis, the whole brain was parcellated into 132 anatomical ROIs based on the Swanson Atlas ^64^. The whole-brain RSFC matrix was computed by Pearson correlations of regionally averaged rsfMRI time courses between pairwise ROIs.

## Acknowledgments

We thank Dr. Yuncong Ma and Ms. Xiaoai Chen for their technical support, and Dr. Xiao Liu for scientific discussion. The present study was partially supported by National Institute of Neurological Disorders and Stroke (R01NS085200), National Institute of Mental Health (RF1MH114224), and National Institute of General Medical Sciences (R01GM141792). The content is solely the responsibility of the authors and does not necessarily represent the official views of the National Institutes of Health.

## Supplementary Figures

**Figure 1 – Figure Supplement 1.**
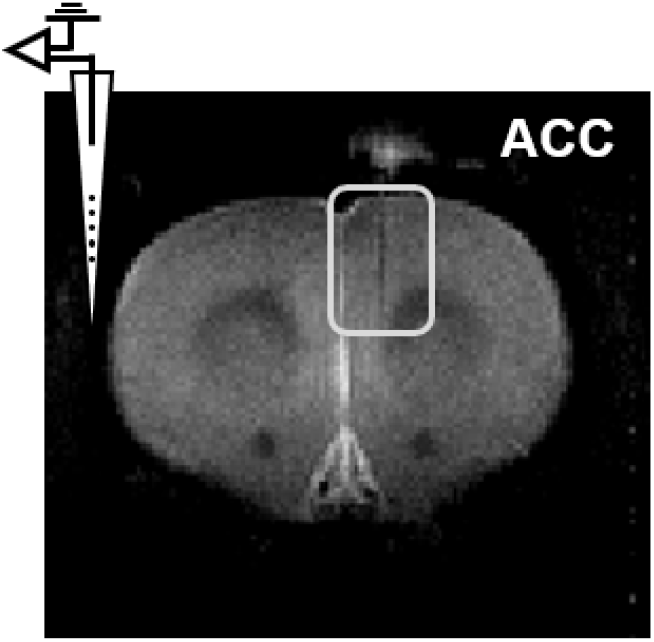
Representative image confirming the electrode location in the anterior cingulate cortex.

**Figure 2 – Figure Supplement 1.**
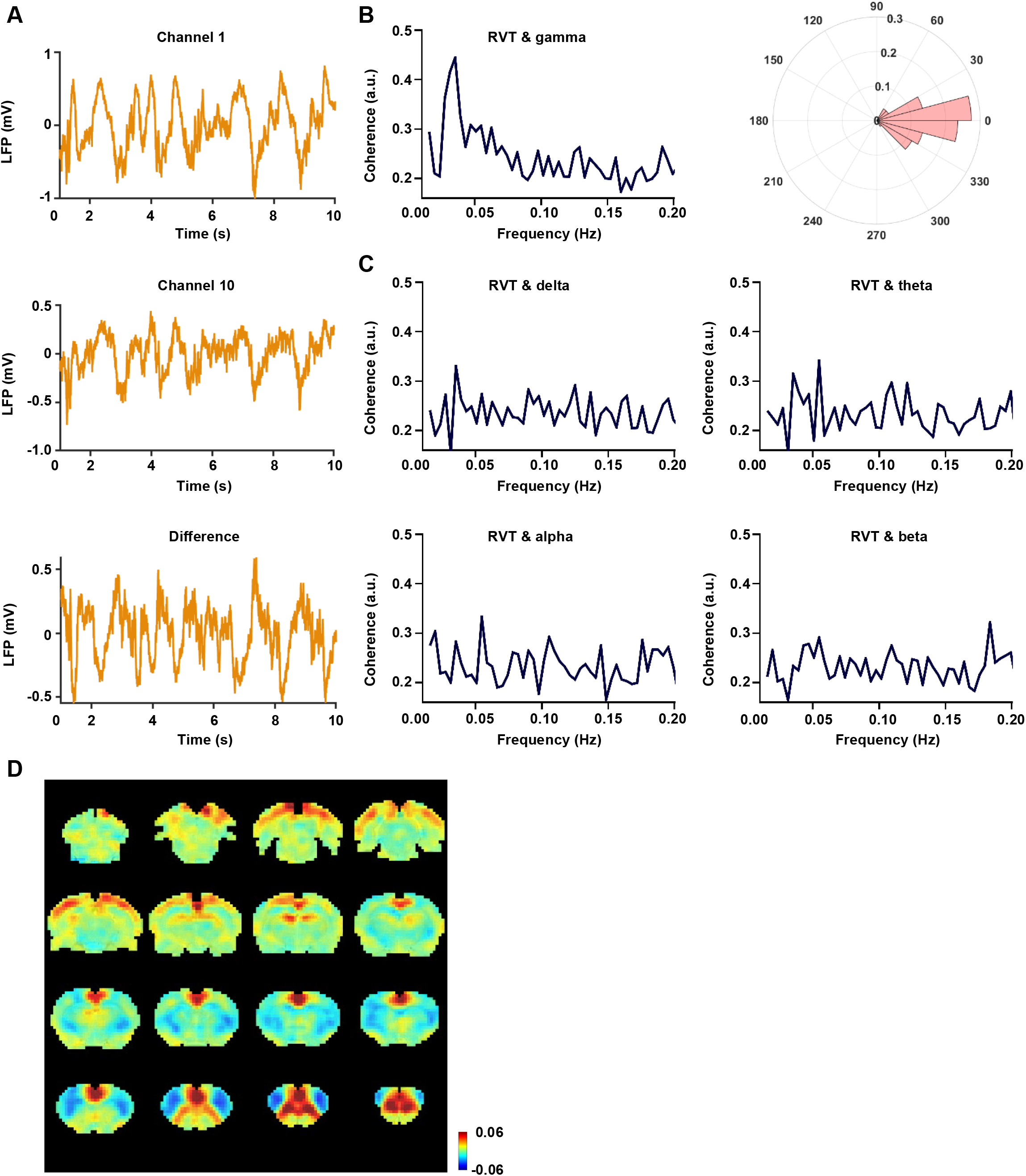
Electrophysiology results obtained using the differential subtraction method. **A**. Exemplar electrophysiology signals from two separate channels: channel 1 (top) and channel 10 (middle), as well as the subtracted signal between the two channels (bottom). Results in panels B-D are all based on the subtracted electrophysiology signal. **B**. Left: coherence between the RVT and gamma-band power (40-100 Hz). Right: phase lag between the RVT and gamma-band power. **C**. Coherence between the RVT and delta band power (1-4 Hz), theta band power (4-7 Hz), alpha band power (7-13 Hz) ans beta band power (13-30 Hz), respectively. **D**. Gamma-band power-derived rsfMRI correlation map.

**Figure 4 – Figure Supplement 1.**
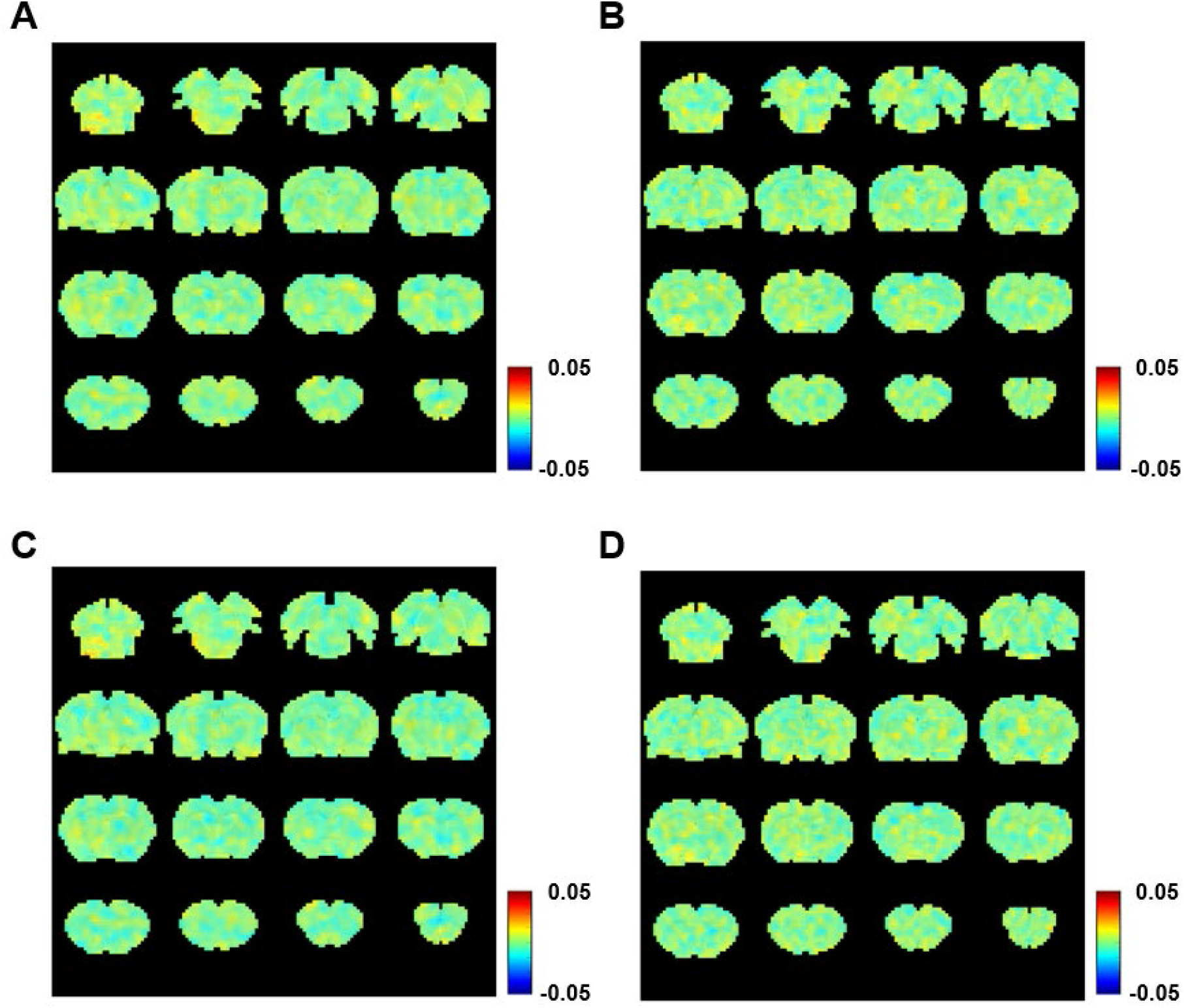
Voxel-wise correlations between the rsfMRI signal and RETROICOR regressor with the regression of the white matter and ventricle signals at (A) light sedation and (B) isoelectric state. Voxel-wise correlations between the rsfMRI signal and RETROICOR regressor without the regression of the white matter and ventricle signals at (C) light sedation and (D) isoelectric state.

**Figure 4 – Figure Supplement 2.**
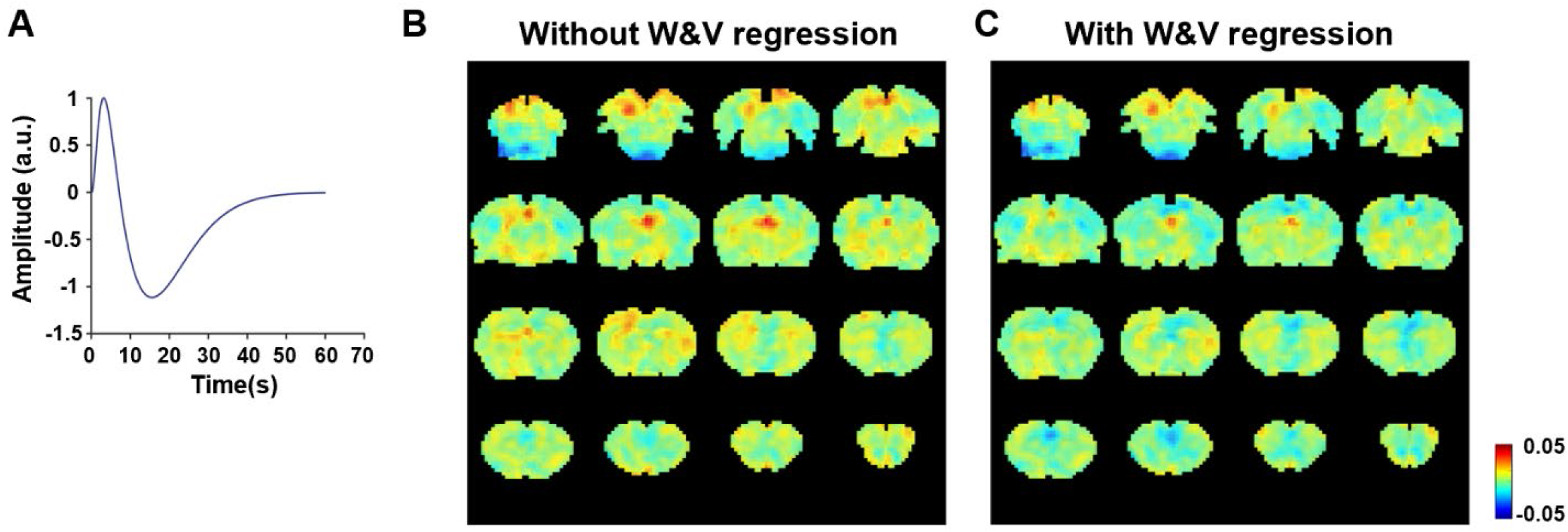
Voxel-wise correlations between the estimated time course of respirational artifacts and rsfMRI signals. **A**. Respiratory response function. Voxel-wise correlations between the estimated time course of respirational artifacts and rsfMRI signals without (**B**) and with (**C**) the regression of the white matter and ventricle signals.

**Figure 3 – Figure Supplement 1.**
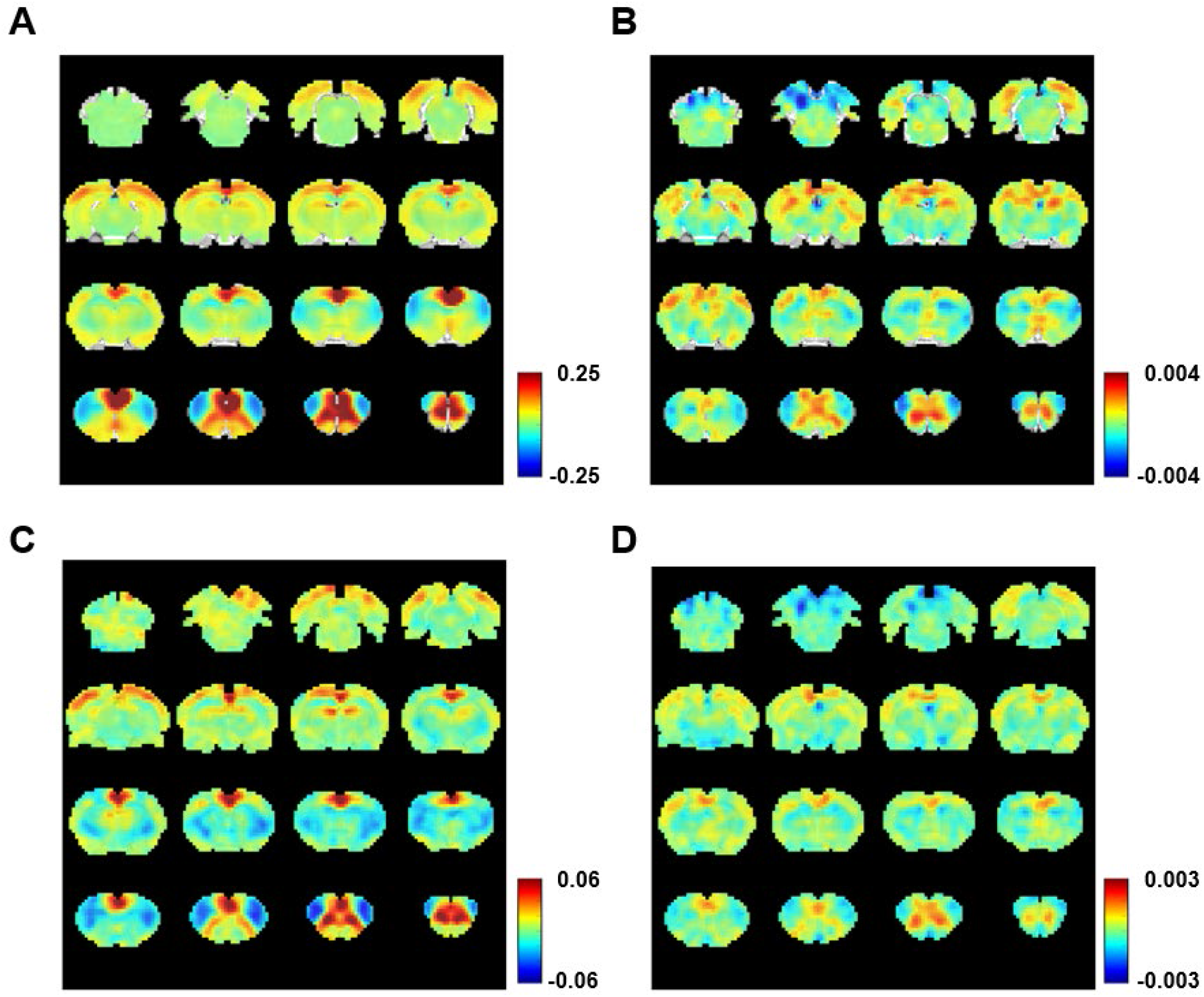
Difference of RSFC maps before and after RVT regression. **A**. Right ACC seedmap after RVT regression; **B**. Difference of right ACC seedmap before and after RVT regression; **C**. Gamma band power-derived correlation map after RVT regression; **D**. Difference of gamma band power-derived correlation map before and after RVT regression.

**Figure 1 – Figure Supplement 2.**
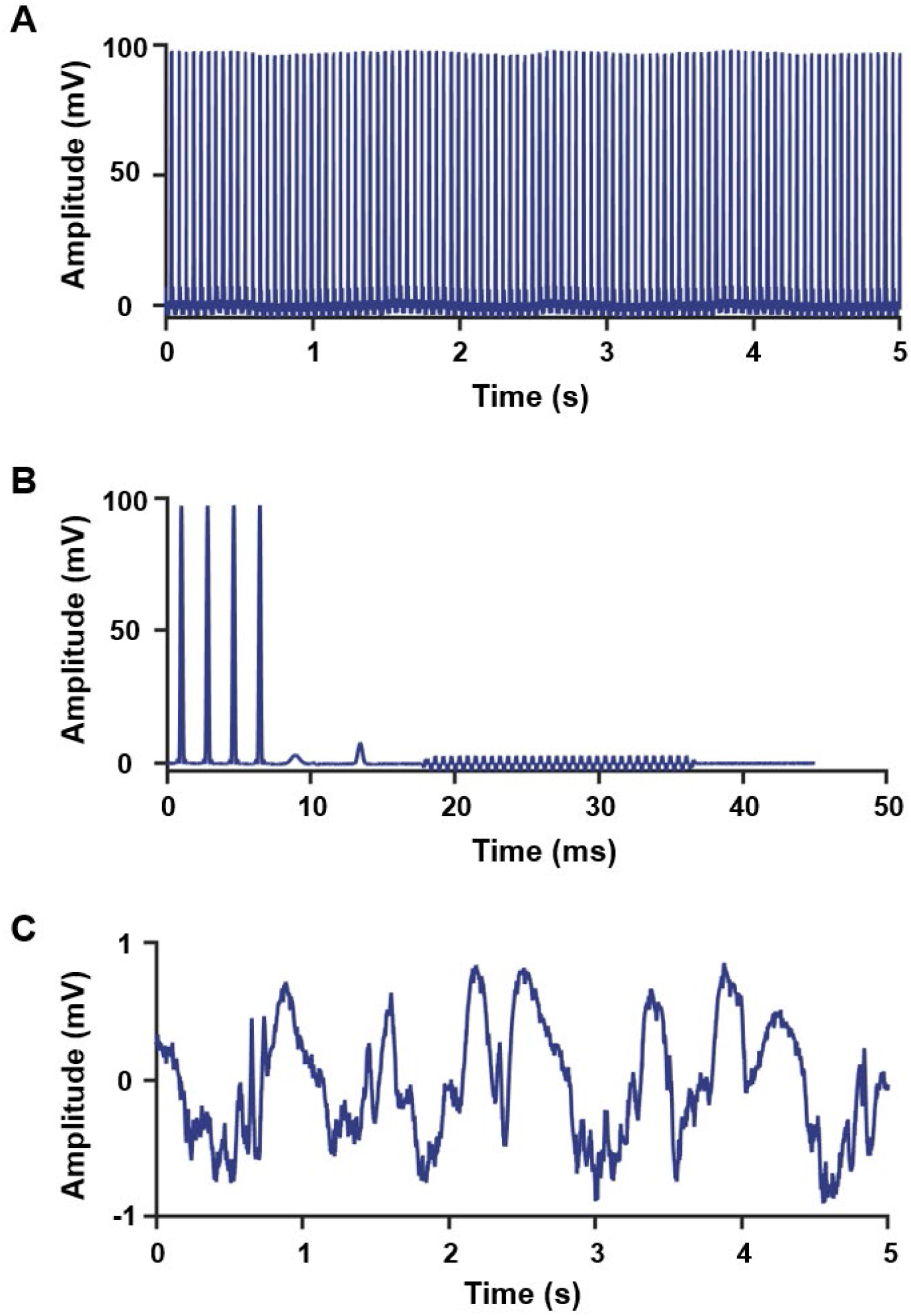
Removal of MRI artifacts from the electrophysiology signal. **A**. an example of raw electrophysiology signals before the denoising of MRI artifacts; **B**. an example of the MRI artifact template. **C**. LFP after MRI artifact denoising.

## Notes

**Conflict of interest:** none.

### Competing Interest Statement

The authors have declared no competing interest.

## References

1 Biswal, B., Yetkin, F. Z., Haughton, V. M. & Hyde, J. S. Functional connectivity in the motor cortex of resting human brain using echo-planar MRI. Magn Reson Med 34, 537–541, doi:10.1002/mrm.1910340409 (1995).

2 Fox, M. D. & Raichle, M. E. Spontaneous fluctuations in brain activity observed with functional magnetic resonance imaging. Nat Rev Neurosci 8, 700–711, doi:10.1038/nrn2201 (2007).

3 Smith, S. M. et al. Correspondence of the brain’s functional architecture during activation and rest. Proc Natl Acad Sci U S A 106, 13040–13045, doi:10.1073/pnas.0905267106 (2009).

4 Murphy, K., Birn, R. M. & Bandettini, P. A. Resting-state fMRI confounds and cleanup. Neuroimage 80, 349–359, doi:10.1016/j.neuroimage.2013.04.001 (2013).

5 Liu, T. T. Noise contributions to the fMRI signal: An overview. Neuroimage 143, 141–151, doi:10.1016/j.neuroimage.2016.09.008 (2016).

6 Caballero-Gaudes, C. & Reynolds, R. C. Methods for cleaning the BOLD fMRI signal. Neuroimage 154, 128–149, doi:10.1016/j.neuroimage.2016.12.018 (2017).

7 Chen, J. E. et al. Resting-state “physiological networks”. Neuroimage 213, 116707, doi:10.1016/j.neuroimage.2020.116707 (2020).

8 Thomason, M. E., Burrows, B. E., Gabrieli, J. D. & Glover, G. H. Breath holding reveals differences in fMRI BOLD signal in children and adults. Neuroimage 25, 824–837, doi:10.1016/j.neuroimage.2004.12.026 (2005).

9 Abbott, D. F., Opdam, H. I., Briellmann, R. S. & Jackson, G. D. Brief breath holding may confound functional magnetic resonance imaging studies. Hum Brain Mapp 24, 284–290, doi:10.1002/hbm.20086 (2005).

10 Birn, R. M., Diamond, J. B., Smith, M. A. & Bandettini, P. A. Separating respiratory-variation-related fluctuations from neuronal-activity-related fluctuations in fMRI. Neuroimage 31, 1536–1548, doi:10.1016/j.neuroimage.2006.02.048 (2006).

11 Wise, R. G., Ide, K., Poulin, M. J. & Tracey, I. Resting fluctuations in arterial carbon dioxide induce significant low frequency variations in BOLD signal. Neuroimage 21, 1652–1664, doi:10.1016/j.neuroimage.2003.11.025 (2004).

12 Chang, C. & Glover, G. H. Relationship between respiration, end-tidal CO2, and BOLD signals in resting-state fMRI. Neuroimage 47, 1381–1393, doi:10.1016/j.neuroimage.2009.04.048 (2009).

13 Hoiland, R. L., Bain, A. R., Rieger, M. G., Bailey, D. M. & Ainslie, P. N. Hypoxemia, oxygen content, and the regulation of cerebral blood flow. Am J Physiol Regul Integr Comp Physiol 310, R398–413, doi:10.1152/ajpregu.00270.2015 (2016).

14 Liu, P. et al. Cerebrovascular reactivity mapping without gas challenges. Neuroimage 146, 320–326, doi:10.1016/j.neuroimage.2016.11.054 (2017).

15 Duyn, J. H., Ozbay, P. S., Chang, C. & Picchioni, D. Physiological changes in sleep that affect fMRI inference. Curr Opin Behav Sci 33, 42–50, doi:10.1016/j.cobeha.2019.12.007 (2020).

16 Picchioni, D. et al. Autonomic arousals contribute to brain fluid pulsations during sleep. Neuroimage 249, 118888, doi:10.1016/j.neuroimage.2022.118888 (2022).

17 Glover, G. H., Li, T. Q. & Ress, D. Image-based method for retrospective correction of physiological motion effects in fMRI: RETROICOR. Magn Reson Med 44, 162–167, doi:10.1002/1522-2594(200007)44:1 (2000).

18 Raj, D., Paley, D. P., Anderson, A. W., Kennan, R. P. & Gore, J. C. A model for susceptibility artefacts from respiration in functional echo-planar magnetic resonance imaging. Phys Med Biol 45, 3809–3820, doi:10.1088/0031-9155/45/12/321 (2000).

19 Windischberger, C. et al. On the origin of respiratory artifacts in BOLD-EPI of the human brain. Magn Reson Imaging 20, 575–582, doi:10.1016/s0730-725x(02)00563-5 (2002).

20 Yuan, H., Zotev, V., Phillips, R. & Bodurka, J. Correlated slow fluctuations in respiration, EEG, and BOLD fMRI. Neuroimage 79, 81–93, doi:10.1016/j.neuroimage.2013.04.068 (2013).

21 Shams, S., LeVan, P. & Chen, J. J. The neuronal associations of respiratory-volume variability in the resting state. Neuroimage 230, 117783, doi:10.1016/j.neuroimage.2021.117783 (2021).

22 Tort, A. B. L., Brankack, J. & Draguhn, A. Respiration-Entrained Brain Rhythms Are Global but Often Overlooked. Trends Neurosci 41, 186–197, doi:10.1016/j.tins.2018.01.007 (2018).

23 Kay, L. M. et al. Olfactory oscillations: the what, how and what for. Trends Neurosci 32, 207–214, doi:10.1016/j.tins.2008.11.008 (2009).

24 Adrian, E. D. Olfactory reactions in the brain of the hedgehog. J Physiol 100, 459–473, doi:10.1113/jphysiol.1942.sp003955 (1942).

25 Fontanini, A., Spano, P. & Bower, J. M. Ketamine-xylazine-induced slow (< 1.5 Hz) oscillations in the rat piriform (olfactory) cortex are functionally correlated with respiration. J Neurosci 23, 7993–8001 (2003).

26 Ito, J. et al. Whisker barrel cortex delta oscillations and gamma power in the awake mouse are linked to respiration. Nat Commun 5, 3572, doi:10.1038/ncomms4572 (2014).

27 Yanovsky, Y., Ciatipis, M., Draguhn, A., Tort, A. B. & Brankack, J. Slow oscillations in the mouse hippocampus entrained by nasal respiration. J Neurosci 34, 5949–5964, doi:10.1523/JNEUROSCI.5287-13.2014 (2014).

28 Biskamp, J., Bartos, M. & Sauer, J. F. Organization of prefrontal network activity by respiration-related oscillations. Sci Rep 7, 45508, doi:10.1038/srep45508 (2017).

29 Zelano, C. et al. Nasal Respiration Entrains Human Limbic Oscillations and Modulates Cognitive Function. J Neurosci 36, 12448–12467, doi:10.1523/JNEUROSCI.2586-16.2016 (2016).

30 Yackle, K. et al. Breathing control center neurons that promote arousal in mice. Science 355, 1411–1415, doi:10.1126/science.aai7984 (2017).

31 Shea, S. A. Behavioural and arousal-related influences on breathing in humans. Exp Physiol 81, 1–26, doi:10.1113/expphysiol.1996.sp003911 (1996).

32 Homma, I. & Masaoka, Y. Breathing rhythms and emotions. Exp Physiol 93, 1011–1021, doi:10.1113/expphysiol.2008.042424 (2008).

33 Folschweiller, S. & Sauer, J. F. Respiration-Driven Brain Oscillations in Emotional Cognition. Front Neural Circuits 15, 761812, doi:10.3389/fncir.2021.761812 (2021).

34 Liotti, M. et al. Brain responses associated with consciousness of breathlessness (air hunger). Proc Natl Acad Sci U S A 98, 2035–2040, doi:10.1073/pnas.98.4.2035 (2001).

35 Lu, H. et al. Rat brains also have a default mode network. Proc Natl Acad Sci U S A 109, 3979–3984, doi:10.1073/pnas.1200506109 (2012).

36 Tu, W., Ma, Z., Ma, Y., Dopfel, D. & Zhang, N. Suppressing Anterior Cingulate Cortex Modulates Default Mode Network and Behavior in Awake Rats. Cereb Cortex 31, 312–323, doi:10.1093/cercor/bhaa227 (2021).

37 Logothetis, N. K., Pauls, J., Augath, M., Trinath, T. & Oeltermann, A. Neurophysiological investigation of the basis of the fMRI signal. Nature 412, 150–157, doi:10.1038/35084005 (2001).

38 Pan, W. J. et al. Broadband local field potentials correlate with spontaneous fluctuations in functional magnetic resonance imaging signals in the rat somatosensory cortex under isoflurane anesthesia. Brain Connect 1, 119–131, doi:10.1089/brain.2011.0014 (2011).

39 Lu, H. et al. Low-but Not High-Frequency LFP Correlates with Spontaneous BOLD Fluctuations in Rat Whisker Barrel Cortex. Cereb Cortex 26, 683–694, doi:10.1093/cercor/bhu248 (2016).

40 Zhang, X., Pan, W. J. & Keilholz, S. D. The relationship between BOLD and neural activity arises from temporally sparse events. Neuroimage 207, 116390, doi:10.1016/j.neuroimage.2019.116390 (2020).

41 Parabucki, A. & Lampl, I. Volume Conduction Coupling of Whisker-Evoked Cortical LFP in the Mouse Olfactory Bulb. Cell Rep 21, 919–925, doi:10.1016/j.celrep.2017.09.094 (2017).

42 Winder, A. T., Echagarruga, C., Zhang, Q. & Drew, P. J. Weak correlations between hemodynamic signals and ongoing neural activity during the resting state. Nat Neurosci 20, 1761–1769, doi:10.1038/s41593-017-0007-y (2017).

43 Liang, Z., Ma, Y., Watson, G. D. R. & Zhang, N. Simultaneous GCaMP6-based fiber photometry and fMRI in rats. J Neurosci Methods 289, 31–38, doi:10.1016/j.jneumeth.2017.07.002 (2017).

44 Du, F. et al. Tightly coupled brain activity and cerebral ATP metabolic rate. Proc Natl Acad Sci U S A 105, 6409–6414, doi:10.1073/pnas.0710766105 (2008).

45 Zhang, Q., Gheres, K. W. & Drew, P. J. Origins of 1/f-like tissue oxygenation fluctuations in the murine cortex. PLoS Biol 19, e3001298, doi:10.1371/journal.pbio.3001298 (2021).

46 Pais-Roldan, P., Biswal, B., Scheffler, K. & Yu, X. Identifying Respiration-Related Aliasing Artifacts in the Rodent Resting-State fMRI. Front Neurosci 12, 788, doi:10.3389/fnins.2018.00788 (2018).

47 Zhao, X., Bodurka, J., Jesmanowicz, A. & Li, S. J. B(0)-fluctuation-induced temporal variation in EPI image series due to the disturbance of steady-state free precession. Magn Reson Med 44, 758–765, doi:10.1002/1522-2594(200011)44:5<758::aid-mrm14>3.0.co;2-g (2000).

48 Birn, R. M., Smith, M. A., Jones, T. B. & Bandettini, P. A. The respiration response function: the temporal dynamics of fMRI signal fluctuations related to changes in respiration. Neuroimage 40, 644–654, doi:10.1016/j.neuroimage.2007.11.059 (2008).

49 Lockmann, A. L., Laplagne, D. A., Leao, R. N. & Tort, A. B. A Respiration-Coupled Rhythm in the Rat Hippocampus Independent of Theta and Slow Oscillations. J Neurosci 36, 5338–5352, doi:10.1523/JNEUROSCI.3452-15.2016 (2016).

50 Zhong, W. et al. Selective entrainment of gamma subbands by different slow network oscillations. Proc Natl Acad Sci U S A 114, 4519–4524, doi:10.1073/pnas.1617249114 (2017).

51 Rojas-Libano, D. & Kay, L. M. Olfactory system gamma oscillations: the physiological dissection of a cognitive neural system. Cogn Neurodyn 2, 179–194, doi:10.1007/s11571-008-9053-1 (2008).

52 Fontanini, A. & Bower, J. M. Variable coupling between olfactory system activity and respiration in ketamine/xylazine anesthetized rats. J Neurophysiol 93, 3573–3581, doi:10.1152/jn.01320.2004 (2005).

53 Herrero, J. L., Khuvis, S., Yeagle, E., Cerf, M. & Mehta, A. D. Breathing above the brain stem: volitional control and attentional modulation in humans. J Neurophysiol 119, 145–159, doi:10.1152/jn.00551.2017 (2018).

54 Schmahmann, J. D. P. D. N. Fiber pathways of the brain. (Oxford University Press, 2009).

55 Illig, K. R. Projections from orbitofrontal cortex to anterior piriform cortex in the rat suggest a role in olfactory information processing. J Comp Neurol 488, 224–231, doi:10.1002/cne.20595 (2005).

56 Garcia-Cabezas, M. A. & Barbas, H. A direct anterior cingulate pathway to the primate primary olfactory cortex may control attention to olfaction. Brain Struct Funct 219, 1735–1754, doi:10.1007/s00429-013-0598-3 (2014).

57 Carmichael, S. T., Clugnet, M. C. & Price, J. L. Central olfactory connections in the macaque monkey. J Comp Neurol 346, 403–434, doi:10.1002/cne.903460306 (1994).

58 Fan, J. et al. Spontaneous brain activity relates to autonomic arousal. J Neurosci 32, 11176–11186, doi:10.1523/JNEUROSCI.1172-12.2012 (2012).

59 Lecrux, C. & Hamel, E. Neuronal networks and mediators of cortical neurovascular coupling responses in normal and altered brain states. Philos Trans R Soc Lond B Biol Sci 371, doi:10.1098/rstb.2015.0350 (2016).

60 Melnychuk, M. C. et al. Coupling of respiration and attention via the locus coeruleus: Effects of meditation and pranayama. Psychophysiology 55, e13091, doi:10.1111/psyp.13091 (2018).

61 Grandjean, J., Schroeter, A., Batata, I. & Rudin, M. Optimization of anesthesia protocol for resting-state fMRI in mice based on differential effects of anesthetics on functional connectivity patterns. Neuroimage 102 Pt 2, 838–847, doi:10.1016/j.neuroimage.2014.08.043 (2014).

62 Tu, W., Ma, Z. & Zhang, N. Brain network reorganization after targeted attack at a hub region. Neuroimage 237, 118219, doi:10.1016/j.neuroimage.2021.118219 (2021).

63 Magri, C., Schridde, U., Murayama, Y., Panzeri, S. & Logothetis, N. K. The amplitude and timing of the BOLD signal reflects the relationship between local field potential power at different frequencies. J Neurosci 32, 1395–1407, doi:10.1523/JNEUROSCI.3985-11.2012 (2012).

64 Swanson, L. W. Brain maps : structure of the rat brain. (Elsevier, 2004).

